# Pinpointing protein structures over broad temperature range using hydrophobic protection

**DOI:** 10.1101/2025.07.08.662548

**Authors:** Fernando de Sá Ribeiro, Luís Maurício T. R. Lima

## Abstract

Protein interactions are dynamic processes influenced by chemical and physical variables. While variable temperatures play an important role in understanding biological processes, their use in protein crystallography has been limited due to the loss of diffraction power. Here, we report the selection of a cryo-protectant that enables X-ray diffraction data collection across a continuum of temperatures, from cryogenic to room temperature. Although several hydrophobic materials effectively preserved samples at 100 K, only a few compounds allowed data collection at 300 K. We identified a hydrophobic grease suitable for both home-source and synchrotron applications, supporting ultrafast and slow diffraction, with exposure times as long as 60 seconds per image at atomic resolution. Data collected between 100 K and 300 K showed no significant effect of exposure time (ranging from 70 msec to 1 second) but revealed an exponential temperature dependence on the overall B-factor. These findings highlight the importance of hydrophobic grease in protecting against environmental variables, enabling optimized data collection across an ultra-wide temperature range.

**Graphical Abstract:** 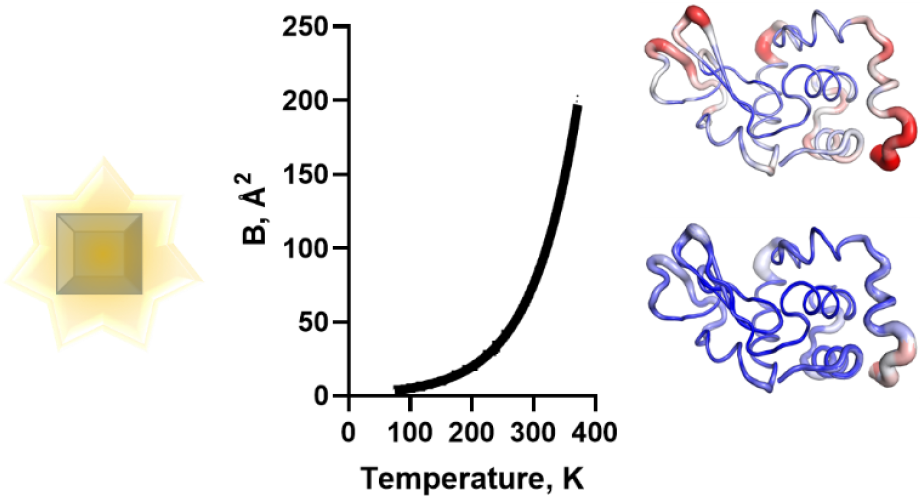

## 1. Introduction

Water is a noble molecule in life ^1^, and great breakthroughs in structural biology came upon learning how to control its properties.

In protein crystallography, the key event opening the road for its development was made in the 1930’s by Prof. John Desmond Bernal and Prof. Dorothy Crowfoot Hodgkin, with their landmark observation that crystals should be protected against dehydration to maintain order and diffraction ^2–4^.

Another major development was the achievement of diffraction under subzero temperatures, denominated cryo-crystallography, paved by the demonstration by Prof. Gregory Petsko in 1975 that replacing or doping the mother liquor with organic compounds resulted in water vitrification, allowing protein single crystal X-ray diffraction at subzero temperatures ^5^ in the absence of ice, and data collection across a broad temperature range (220 K – 300 K) ^6^. Later in 1992, Prof. Petsko and his group further used this solvent replacement approach to demonstrated that an apparent biphasic transition occurs in RNAse crystal diffraction over a temperature transition from cryo (98 K) to high (320 K) temperature ^7,8^, using different co-solvent (2-methyl-2,4-pentanodiol; methanol) and crystal mounting (capillary; glass fiber) strategies at each temperature. They explained the apparent biphasic transition observed in the Debye-Waller factor as a function of temperature seen in their work ^6–8^ to being dominated by an harmonic motion at higher temperatures progressively decreasing and dominated by harmonic vibration of individual atoms at temperatures below 200 K.

The B-factor, or temperature factor, is related to the mean-square atomic displacement (*X*^2^) by the Debye-Waller function:

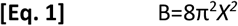

The *X*^2^ comprise contribution from atomic fluctuation (*X*^2^ _A_), conformational motions (*X*^2^ _C_), rigid body vibration (*X*^2^_RB_ ) and crystal lattice disorders (*X*^2^_LT_ ) ^9^, which can be dissected by exploring physical variables affecting the B-factor. However, while the crystal structure is highly reproducible ^10,11^, the B-factor vary largely between crystals, instruments, and co-solvents, limiting the use of multiple single crystals in the investigation of effects of variables in the B-factor or in serial crystallography.

The reproducibility of B-factor was finally achieved by mathematically normalization ^12,13^:

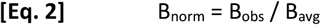

where B_norm_ is the normalized B-factor, B_avg_ is the average B-factor and B_obs_ is the observed F-factor. While normalization allow several analysis including the correlations between B-factor and structural changes, and can be easily implemented in data processing pipelines in structural biology methods as such as single crystal and serial crystallography, cryo-EM ^14,15^ and others ^16,17^ (exporting PDB with normalized B-factor), it does not allowed interpreting the absolute Debye-Waller factor.

The experimental reproducibility in B-factor could be achieved by crystal stabilization using hydrocarbon grease ^18,19^, allowing demonstration of the dominance of whole-system rigid-body motion on the temperature-dependence B-factor change, well described by a single-exponential dependence of the B-factor as a function of temperature for all atoms of the system (protein / non-protein) with a thermal constant “k” of about 0.004 K^-1^ as follow:

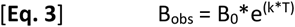

where B_0_ is the B-factor extrapolated to zero Kelvin from regression (about 5 Å^2^), k is the thermal constant, and T is the data collection temperature (in Kelvin).

After the advances obtained with cryo-crystallography, a recent revival of the room temperature data collection returned to spotlight motivated by serial crystallography, and alternative non-dehydration techniques have been pursued such as tubbing (capillaries or other polymer devices) ^20^, glues ^7,21^, silicone ^22^, lipids ^23^, oils ^24–26^, mineral oil-based grease ^27^ vegetable ^28^ and animal fat ^29,30^. A recent work on the X-ray diffraction at high resolution (1.40 to 1.68 Å) of tetragonal Proteinase-K from room temperature (293 K) to extreme high temperatures (363 K, 90 °C), conducted with crystals embedded into Paratone-N oil as originally proposed by Hope in 1988 ^24^, suggested a progressive increase in the Wilson B-factor (from 23 to 34 Å^2^) as a function of temperature ^31^. Unfortunately, this work did not addressed temperatures below 20 °C.

The comparative use of these crystal stabilization approaches for broad-range temperature measurements without changes in crystal composition have not been tested thoroughly, and there is a large gap between cryogenic and room temperatures in the literature and the PDB database that still need to be explored (Fig. 1).

**Figure 1.**
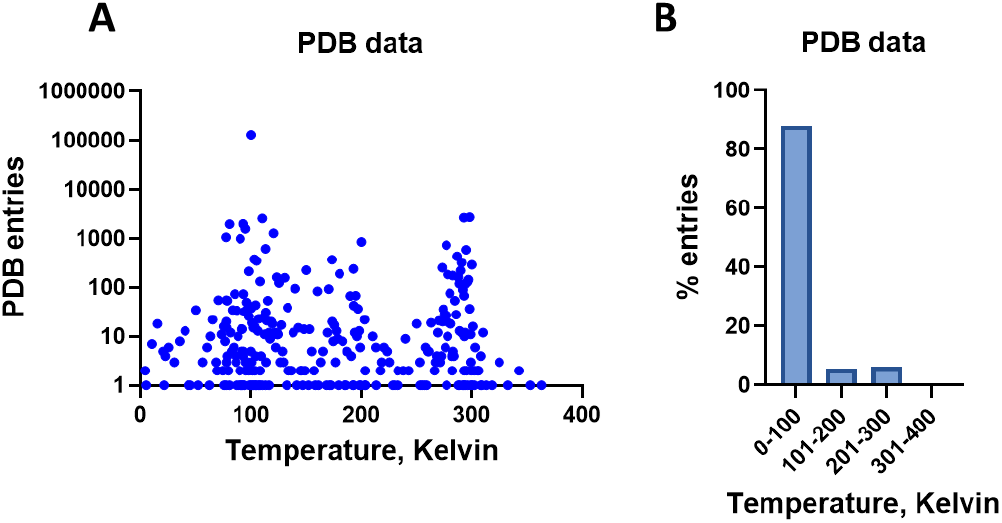
Distribution of reported X-ray diffraction temperature. Data was retrieved from the RCSB (access May, 2025) A) All structures deposited in the PDB by temperature (notice log scale). B) All structures deposited in the PDB by temperature range (100K interval).

Several limitations still hold at present, such as study of continuous temperature dependence over broad temperature, from cryo to above room temperature, allowing exploring the temperature effect on conformational changes, ligand interaction, raw B-factor, as well as conducting of measurements at long exposure in home source or synchrotron without major radiation damage.

In this study we explore a large set of potential protective agents for data collection at cryogenic and ambient conditions, and explore the protection against high synchrotron flux over a broad temperature range and the application in the interpretation of conformational and B-factors changes.

## 2. Material and Methods

### 2.1. Material

Hen egg white lysozyme (UNIPROT ID P00698) was obtained from Fluka (Cat #62971; Lot #BCG4805V) and kept at -20 °C until use. Apiezon N and Apiezon T were obtained from Sigma-Aldrich. Vaseline (petroleum jelly; Vasenol), mineral oil (Farmax), lard and olive oil were purchased from local stores. Silicone oil (dimethyl siloxane, trimethylsiloxy-terminated; Dow Corning 360 Medical Fluid, 350 CST) was a kind donation from Hygeia Biotech S/A. We did not test Paratone-N oil since we did not have access to this product. All other reagents were of analytical grade.

### 2.2. Methods

#### 2.2.1. Protein crystallography

Crystals of tetragonal (P4_3_2_1_2) hen egg white lysozyme (HEWL) were obtained by vapor diffusion, sitting-drop technique, using Corning 3552 plates with 1 μL protein solution (HEWL 50 mg/mL in water), and 1 μL precipitant composed by 100 mM sodium acetate pH 4.6, 1.7 M NaCl, with (synchrotron data collection) or without (home source data collection) 25 % ethylene glycol, equilibrated against 80 μL precipitant solution at 20 °C ± 2 °C ^32^. Diffraction-quality crystals were used after two days. Crystals were manually picked using 20 μm nylon CryoLoop™ (Hampton Research), soaked in the crystallization solution supplemented or not (in case of having ethylene glycol) with glycerol (for 30% v/v final concentration), following by immersion into the cryoprotection. The crystals were mounted onto the goniometer with N_2g_ stream set at the desired temperature for data collection, and full set was collected as depicted below. For each dataset a new crystal was used.

#### 2.2.2. Data collection and processing

Two workflows were used in the X-ray diffraction and data collection.

a. **Home source** – Crystals were diffracted using CuKα radiation from a D8-Venture diffractometer (Bruker AXS Inc.; installed in the CENABIO-UFRJ) operating at 50 kV and 1.1 mA from a 30 W air-cooled |μS microfocus source (Incoatec), and images were recorded on a Photon II detector (Bruker). Temperature was controlled with N_2g_ stream (flow of 1.2 L/h) set to the desired temperature, using a CryoStream 800 (Oxford Cryogenics). All datasets were collected with frames of 1 min exposure / 0.5° oscillation, reaching at least 99% completeness and 1.5 Å resolution (for Friedel pairs set to true). All data collection, indexing, integration and scaling was performed by *Proteum3* (Bruker AXS Inc.), and further analyzed with *Truncate* – CCP4 ^33^.
b. **Synchrotron** – Data was obtained by diffraction with radiation at 16 keV (0.7749 Å) and collected with Pilatus 2M (Dectris) detector set at 112 mm from crystal for all datasets, allowing a maximum resolution of 0.947 Å. Temperature was controlled by N_2g_ stream at 8 L/min using CryoStream 700 (Oxford Cryogenics). All datasets were collected with 0.5° oscillation, in a total of 180°, resulting in 360 images, processed with XDS implemented in MNCAutoproc ^34,35^ written by Dr. Andrey Nascimento (Manaca beamline, Sirius, CNPEM, Campinas, SP, Brazil) after Autoproc ^36^, set to resolution at CC½ = 50%. Beam attenuation was changed inversely proportional to exposure time per frame (70 msec, 1 sec, 5 sec, 30 sec) in order to decrease flux (photons / sec) resulting in close total photon count per frame. While we used nylon cryoloops for all data collection, tests conducted with commercially available Kapton-based multi-hole also showed good adherence and easy for cleaning up (*data not shown*).

#### 2.2.3. Data processing

All data were further processed with *C*.*C*.*P*.*4*. v7.0.071 ^35^. Structures were solved by molecular replacement with rigid body refinement using RefMac v5.8.0238 ^37 38^ with PDB ID 3A8Z ^39^, followed by restrained refinement in *Refmac* ^37^. Further real space modeling was performed with *C*.*O*.*O*.*T*. v0.8.9.2 ^40^ and an additional restrained refinement with *Refmac*, always in default mode to avoid bias.

Details of the crystal parameters, data collection, and refinement statistics are compiled in Table S1 of the Supporting Information. All molecular representations were created using PyMOL v2.0 (The PyMOL Molecular Graphics System). The final atomic coordinates have been submitted to the Protein Data Bank (PDB), with accession codes provided in the Supporting Information.

#### 2.2.4. Data analysis

Graphics were generated with GraphPad Prism v. 8.0.2 for Windows (GraphPad Software, San Diego, California USA).

## 3. Results

### 3.1. Screening of thermoprotectants

We have tested embedding lysozyme crystals into varying compounds and evaluated the stability for X-ray data diffraction at cryogenic condition (100 K) and room temperature. At 100 K, all tested compounds were satisfactory at keeping data diffraction at 1.5 Å, although resulting in dissimilar data diffraction quality as judged by the data collection parameters (Table 1). However, we observed loss of diffraction power at room temperature for most tested stabilizing agents, except for the hydrocarbon greases vaseline and Apiezon (Table 1).

**Table 1.** Statistical analysis of data collection using varying protective agents. Lysozyme crystals (n=3 per condition; average and standard deviation) were embedded into candidate stabilizing agents and subjected to full X-ray data collection at a target resolution of 1.5 Å.

|  | 100 K |  |  |  |  | 300 K |  |  |  |  |
| --- | --- | --- | --- | --- | --- | --- | --- | --- | --- | --- |
|  | a=b | c | <sup>w</sup>Bf | lowI/σ | highI/σ | a=b | c | <sup>w</sup>Bf | lowI/σ | highI/σ |
| **Apiezon® N** | 78.24 | 37.34 | 10.97 | 109.89 | 9.55 | 79.23 | 38.10 | 14.50 | 35.33 | 2.65 |
|  | ±0.12 | ±0.05 | ±1.04 | ±6.53 | ±1.70 | ±0.06 | ±0.01 | ±1.07 | ±27.02 | ±0.76 |
| **Apiezon® T** | 78.13 | 37.15 | 11.23 | 110.68 | 7.22 | 79.00 | 38.10 | 14.13 | 106.65 | 3.34 |
|  | ±0.20 | ±0.08 | ±1.58 | ±9.69 | ±3.24 | ±0.09 | ±0.10 | ±3.36 | ±31.52 | ±0.42 |
| **Glycerol** | 78.23 | 37.03 | 13.27 | 145.33 | 6.69 |  |  | N/A |  |  |
|  | ±0.30 | ±0.05 | ±1.22 | ±30.35 | ±3.20 |  |  |  |  |  |
| **Lard** | 78.38 | 37.26 | 10.97 | 101.98 | 9.67 |  |  | N/A |  |  |
|  | ±0.09 | ±0.08 | ±0.91 | ±5.85 | ±0.37 |  |  |  |  |  |
| **Mineral Oil** | 78.55 | 37.16 | 10.67 | 190.47 | 14.61 |  |  | N/A |  |  |
|  | ±0.12 | ±0.05 | ±0.11 | ±21.64 | ±1.35 |  |  |  |  |  |
| **No Cryo** | 77.89 | 37.27 | 13.37 | 101.62 | 3.86 |  |  | N/A |  |  |
|  | ±0.27 | ±0.06 | ±0.82 | ±15.25 | ±0.57 |  |  |  |  |  |
| **Olive Oil** | 78.15 | 37.16 | 10.80 | 79.92 | 6.84 |  |  | N/A |  |  |
|  | ±0.19 | ±0.14 | ±0.67 | ±7.82 | ±0.85 |  |  |  |  |  |
| **PEG 400** | 77.23 | 37.42 | 13.03 | 158.00 | 8.42 |  |  | N/A |  |  |
|  | ±0.12 | ±0.06 | ±1.04 | ±20.41 | ±2.56 |  |  |  |  |  |
| **PEG 6000** | 78.53 | 37.16 | 11.17 | 159.96 | 9.18 |  |  | N/A |  |  |
|  | ±0.09 | ±0.01 | ±0.44 | ±11.95 | ±1.58 |  |  |  |  |  |
| **Silicone Oil** | 78.48 | 37.13 | 11.30 | 184.69 | 10.51 |  |  | N/A |  |  |
|  | ±0.03 | ±0.03 | ±0.47 | ±24.83 | ±2.07 |  |  |  |  |  |
| **vaseline** | 78.08 | 37.30 | 12.47 | 99.60 | 6.65 | 79.08 | 38.26 | 14.23 | 92.32 | 3.12 |
|  | ±0.49 | ±0.07 | ±0.71 | ±6.40 | 0.80 | ±0.13 | 0.17 | ±1.64 | ±29.32 | ±0.22 |

a, b and c are cell parameters (Å)
<sup>w</sup>Bf is Wilson B-factor
lowI/σ<sub>Avg</sub> is low-resolution Intensity / sigma
highI/σ is high-resolution Intensity / sigma
N/A – Not available, due to loss of diffraction power.

Although all stabilizing agents allowed full single crystal X-ray data diffraction and structure solving at same target resolution (1.5 Å), they impacted the data quality (Table 1; Fig. 2). A positive correlation between I/σ at high resolution and low resolution was observed (p=0.022), with negative correlation between I/σ at high resolution and the Wilson B-factor (p=0.030) (Fig. 2). Although the higher Intensity with lower Wilson B-factor was obtained with mineral oil, this tested agent did not allow data collection at 300 K. The other better agents under these criteria for data collection at 100 K were silicone oil and Apiezon, although silicone oil also resulted in low of diffraction power at 300 K. The hydrocarbon grease Apiezon N is recommended for a working range of 4 K to 303 K (-269 °C to 30 °C), while the Apiezon T is recommended for temperature ranging from 283 K to 393 K (10 °C to 120 °C).

**Figure 2.**
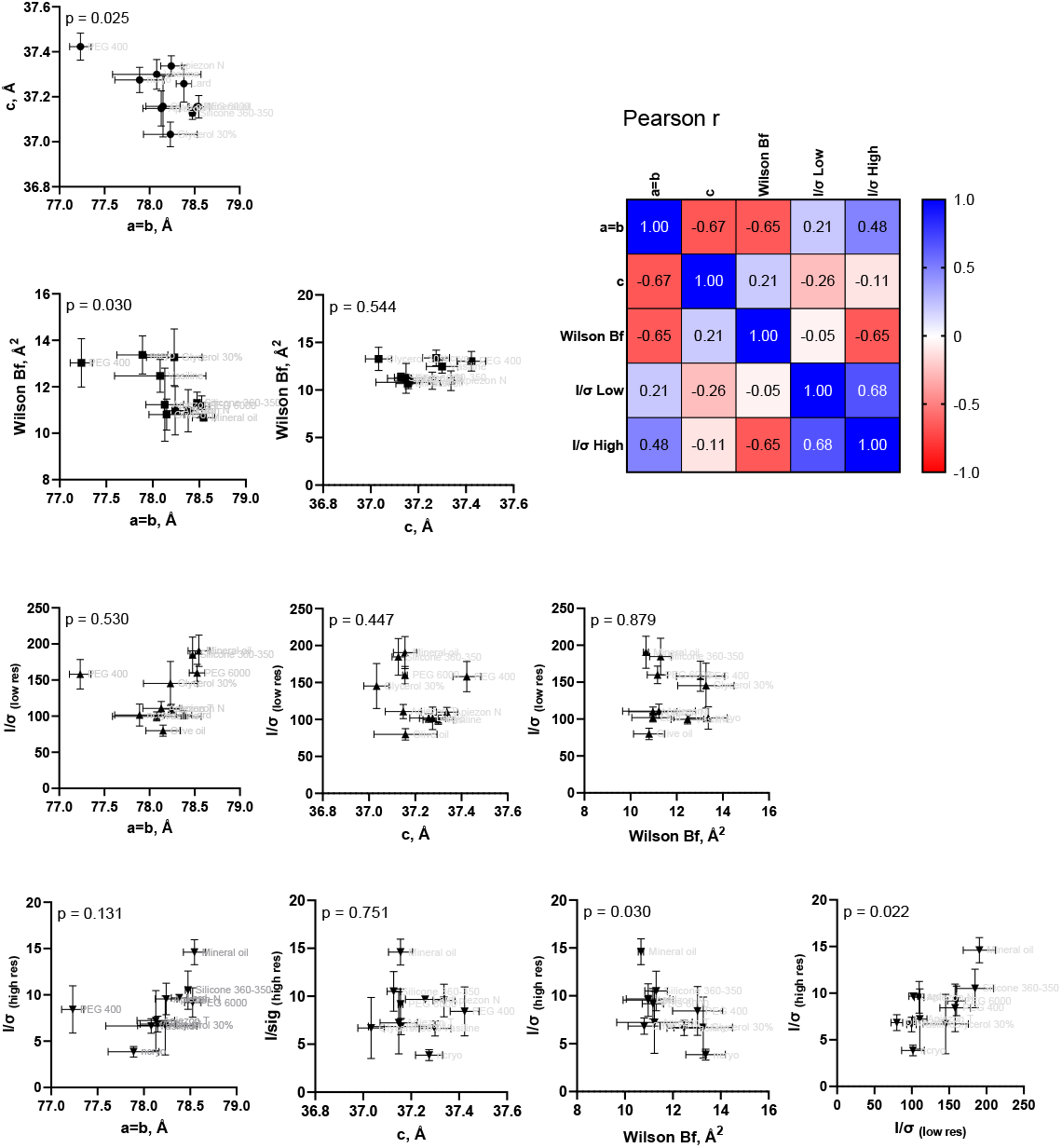
Correlation analysis between data with varying stabilizing agents. Pearson correlation r and p-value were obtained for data obtained from full dataset collected from X-ray single-crystal diffraction at 100 K. Symbols and bars represent average and standard deviation (n=3) respectively.

The structural finds showed a great similarity between all models to both cases (100 K and room temperature) (Fig. 3). The B-factor analysis of the structures reveals overall increase from data collected at 100 K and 300 K for all conditions, which becomes similar in distribution over the polypeptide chain upon normalization (Eq. [2]), indicating no major distinction between them (Fig. 4). Since we aimed to collect data under varying temperatures as continuous variable, from cryo to room temperature, we choose the Apiezon N for further investigations.

**Figure 3.**
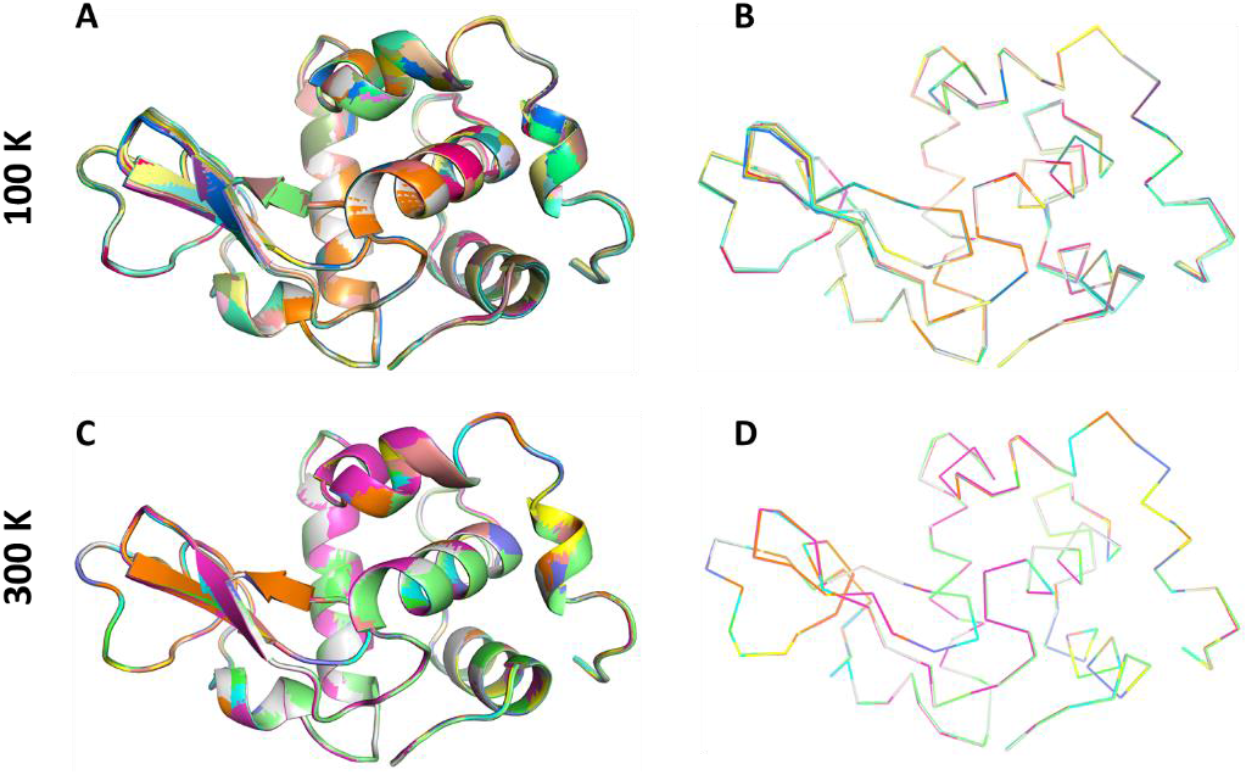
Alignment of crystal structures. Structures solved using home-source X-ray and varying stabilizing agents (Table 1; n=3/compound) were superposed using PyMOL. A) 100 K, carton representation. B) 100K, backbone representation. C) 300K, carton representation. D) 300K, backbone representation.

**Figure 4.**
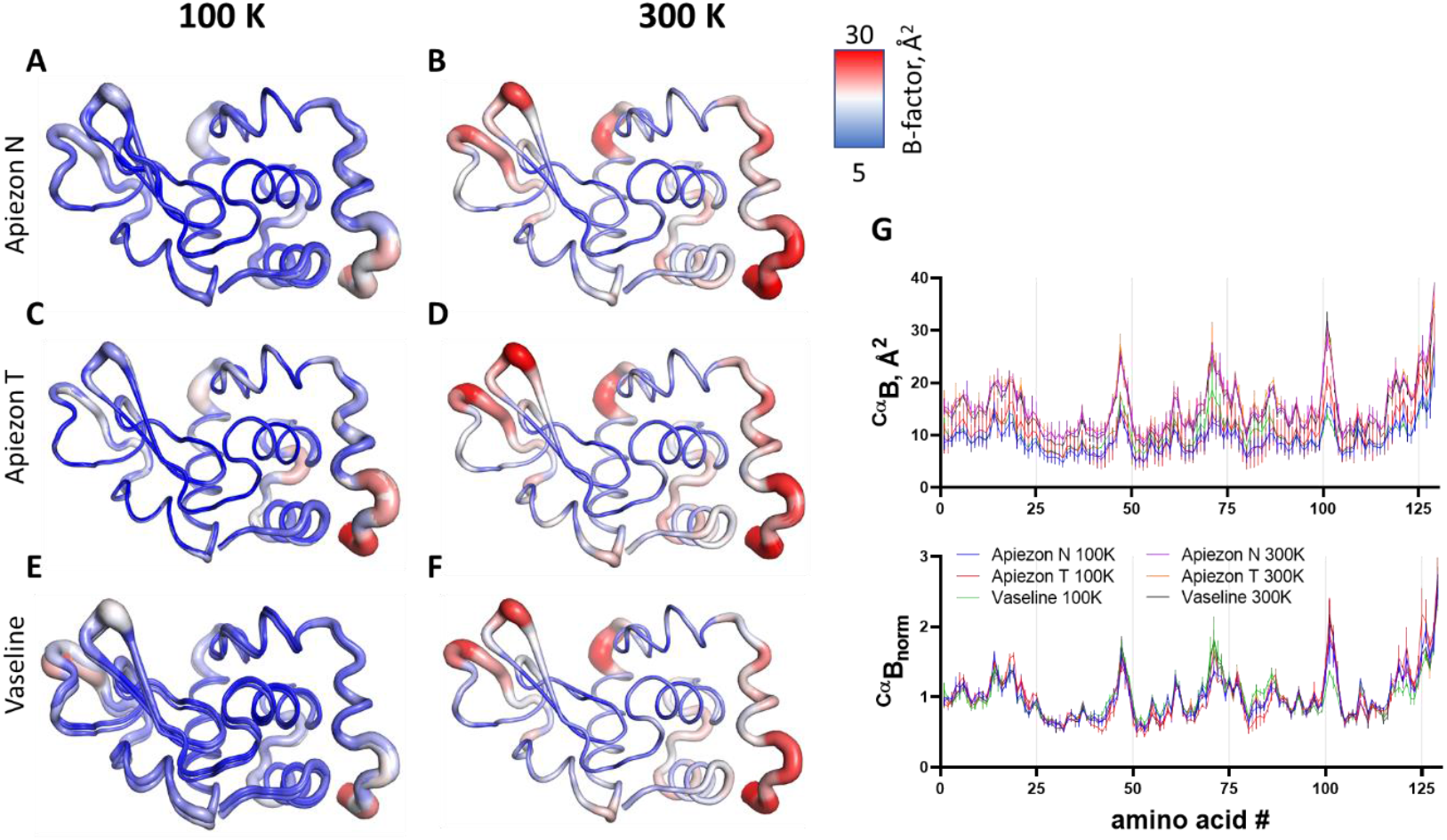
B-factor distribution crystallographic structures. Structures solved from X-ray data diffraction of single crystal (n=3/group) using varying stabilizing agents are represented by B-factor scale according to the presented color scale. A) Apiezon N (100 K); B) Apiezon N (300 K); C) Apiezon T (100 K); D) Apiezon T (300 K); E) vaseline (100 K); F) vaseline (300 K). G) The raw and the normalized B-factors for each structure (average and standard deviation; n=3/condition) are presented.

### 3.2. Grease thermal protection against short- and long –exposure synchrotron radiation

We further tested the crystal quality stability against short and prolonged data collection using synchrotron radiation. Crystal diffraction data collection under 5 sec and 30 sec exposure per frame, attenuated proportionally for similar photon dose, did not affected the resolution, B-factor or cell parameters substantially, despite increase in I/σ in the low resolution without benefits in resolution (Table 2). Thus, the use of hydrocarbon grease could benefit longer exposures for low-diffracting crystals.

**Table 2.** Exposure time-dependence on X-ray diffraction data quality by protein crystals. Full dataset was collected form HEWL crystals embedded into hydrocarbon grease with varying exposure time compensated for transmission for close dose exposure per frame.

| Exp. time, sec | Transmission, % | Flux (ph/s) | Dose / frame | Res., Å (CC<sub>1/2</sub>=0.5) | I/sig low | Wilson Bf | a=b, Å | c, Å |
| --- | --- | --- | --- | --- | --- | --- | --- | --- |
| 0.07 | 100 | 3.56E+11 | 2.49E+10 | 1.02 | 51.39 | 9.10 | 78.96 | 36.75 |
| 1 | 10 | 3.48E+10 | 3.48E+10 | 0.99 | 63.05 | 9.20 | 79.02 | 36.77 |
| 5 | 2 | 6.08E+09 | 3.04E+10 | 0.99 | 71.22 | 8.90 | 79.00 | 36.80 |
| 30 | 0.3 | 8.91E+08 | 2.67E+10 | 1.00 | 74.89 | 8.70 | 79.03 | 36.80 |

### 3.3. Thermal-dependence of crystal quality upon synchrotron radiation X-ray diffraction data collection

Tetragonal lysozyme crystals embedded into the hydrocarbon grease were subjected to full dataset data X-ray diffraction from 100 K to 300 K in 25 K intervals, with varying exposure time (*70 msec and 1000 msec*) and attenuation resulting in similar dose/frame. Data from integration revealed minor variations in cell dimensions (a=b *vs* c; Fig. 5A), which were linearly correlated (Pearson r=0.82, 95% CI = 0.65 to 0.92, p<0.0001) (Fig. 5B).

**Figure 5.**
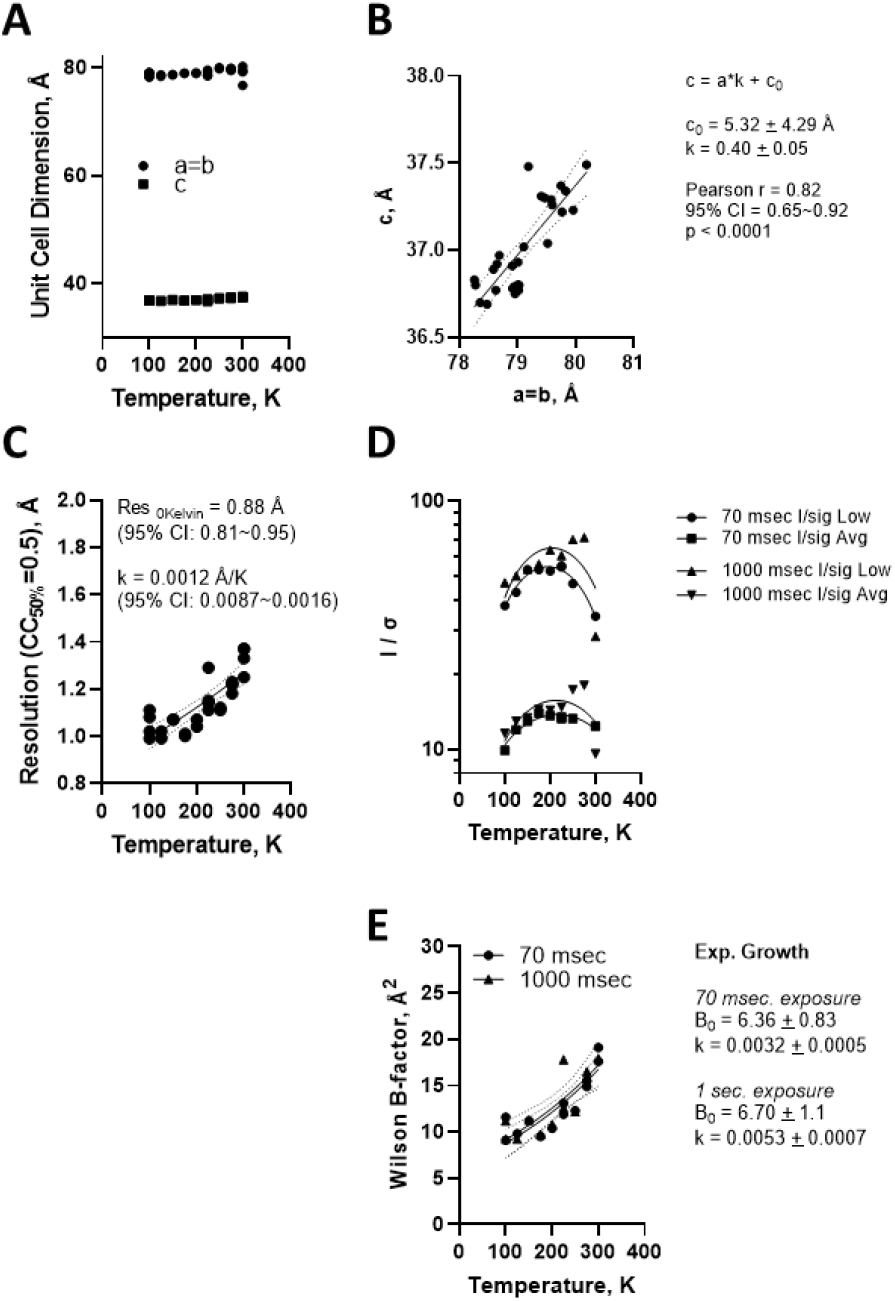
Temperature-dependence effect on X-ray diffraction crystal properties. Apiezon-stabilized crystals were diffracted using synchrotron radiation and data quality were evaluated against temperature for **A)** cell parameters, **B)** correlation between cell parameters, **C)** resolution threshold, **D)** I/sigma and **E)** Wilson B-factor.

The resolution limit, automatically defined by CC_1/2_ = 0.5 as criteria, showed small but progressive decrease from about 1.0 Å at 100 K to about 1.3 Å at 300 K (Fig. 5C).

There occurred a biphasic dependence of the reflection intensity as a function of increasing temperature (Fig. 5D), increasing from 100 K to about 200 K followed by a decrease up to 300 K.

The Wilson B-factor showed an exponential increase as a function of temperature, with overall extrapolated zero-point B-factor ^Wilson^Bf_0_ = 6.27 ± 0.56 Å^2^ (*B*_*0*_ *= 6*.*36 + 0*.*83 for 70 msec, and B*_*0*_ *= 6*.*70* ± *1*.*1 for 1 sec exposure*) and exponential factor k = 0.0034 ± 0.0004 Å^2^/K (*k = 0*.*0032 ± 0*.*0005 for 70 msec, and k = 0*.*0053 ± 0*.*0007 for 1 sec exposure*) (Fig. 5E).

These data indicate the ability for the hydrocarbon grease to support data diffraction continuously over a broad range of temperature using 4^th^ generation synchrotron radiation at both short and long X-ray exposure. Further investigations with other crystals from other proteins are required to understand the benefits of crystal protection in the at the access to diffraction information transition range of 200 K.

## 4. Discussion

Proteins are highly dynamic and can populate a vast conformational space. Characterizing these polymorphs require chemical and physical variables associated to spectroscopic and other state-of-the-art techniques, such as NMR, cryoEM and crystallography. While NMR and cryoEM data collection are performed in orthogonal temperature conditions (above zero Celsius for NMR, and around 77 Kelvin for cryoEM, liquid nitrogen temperature), crystallography could benefit from near-zero temperature ^41^ up to 90 °C ^42^. However, a significant gap persists in crystallography between cryogenic (∼100 K) and room-temperature conditions—a “cryogenic valley of death” largely due to the water phase transition near 200 K. In this work, we evaluated various readily available compounds as stabilizing agents for single-crystal X-ray diffraction data collection across a temperature range from cryogenic to room temperature. Among these, we identify the effectiveness of vaseline and Apiezon—hydrocarbon greases known for their broad temperature stability and low impurity levels. Apiezon N, in particular, offered advantages due to its viscosity, which promotes crystal adherence and prevents dehydration during prolonged exposures (up to hours).

Apiezon greases may have some limitations. Their transparency is lower than that of oils, which may restrict the size of crystals suitable for diffraction. Additionally, while Apiezon is effective for single-crystal diffraction across the tested temperature range, its high viscosity could pose challenges for serial crystallography, requiring further optimization. In this study, we found that animal and vegetable greases were unsuitable for single-crystal X-ray diffraction at room temperature (Table 1), despite prior demonstrations of their applicability in serial crystallography ^28–30^. We were unable to test these compounds for serial crystallography due to lack of access to the technique.

The advent of cryo-crystallography enabled long X-ray exposures with high-resolution diffraction, particularly under high-intensity beamlines ^43^. Hope ^24^ proposed a linear relationship between temperature and the B-factor for crystals embedded in Paratone-N oil:

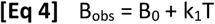

where B_obs_ is the B-factor at temperature T, and B_0_ is the *‘zero-point*’ B-factor (at zero Kelvin), and k to be a linear proportionality constant. Later, Petsko and colleagues ^7^ suggested a biphasic model, with distinct linear relationships below and above 200 Kelvin:

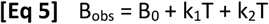

In their review on protein dynamics, Ringe and Petsko ^9^ noted that while mean-square displacement *X*^2^ could be refined with precision, determining their absolute accuracy remained challenging. Reproducible absolute B-factor were only achieved through stabilization with hydrophobic grease ^18,19^. Ringe and Petsko further hypothesized that below 100 K, the B-factor would become temperature-independent and uniform across all atoms, yielding an extrapolated zero point B-factor of 0.4 Å^2 9^.

Contrary to this biphasic model, our data - spanning cryogenic to room temperature (100 K to 325 K) at 1.5 Å resolution - demonstrate that the temperature dependence of the B-factor follows a single-exponential (Eq. [3]). Using this model, we derived a thermal dependence constant (“k”) of about 0.0045 ± 0.0003 K^-1^ and a zero-point B-factor B_0_ of about 5.7 ± 0.5 Å^2^ from home-source diffractometer data (∼ 3 hours data collection time. Synchrotron data (70 m sec to 1 sec exposure/image at 1.0 - 1.3 Å resolution) yielded k = 0.0034 ± 0.0004 K^-1^ and B_0_ = 6.3 ± 0.6 Å^2^. Notably, hydrocarbon grease compatibility was further supported by consistent results across ultrafast (70 msec) and ultra-long (30 sec) synchrotron exposure (Table 2).

Cryo-electron microscopy (cryo-EM) is a technique made possible by vitrification of water through ultrafast cooling ^44^. The significance of B-factor in crystallography is extensive to cryo-EM as originally proposed by Prof. Peter B. Rosenthal and Prof. Richard Henderson ^45^, calculated form the model-independent Guinier plot of the structure factor amplitude (F) against reciprocal resolution. The cryo-EM B-factor incorporates signal attenuation from multiple sources and, despite the near-atomic resolution achieved by the cryo-EM along with other advances in the field such as direct electron detectors, it is not unusual to find overall B-factor close or above 100 Å^2^. Model-generated maps based on B-factors are routinely used ^46^ demonstrating the importance of a proper use and interpretation of this physical parameter. While vitrification can be achieved with pure water and dilute aqueous solutions as originally demonstrated ^47^ and has been adopted as gold-standard in current practice, the use of additives as cryoprotectants as initially proposed by Petsko for protein crystallography ^5^, is still in early stages for cryo-EM ^48^. Further investigations on the origin and the physical meaning of each component of the crystallographic and the cryo-EM B-factors may provide deep understanding on protein motion and dynamics in the vitreous world, from single particle to cellular milieu.

To validate the thermal dependence of the crystallographic B-factor, further studies should test additional proteins. Confirming the universality of the proportionality constant (“k”) and zero-point B-factor B_0_ - both from raw data (Wilson B-factor) and refined atomic models - would clarify molecular motion in the crystalline state. Such verification could also drive broader advances in structural biology. In addition, advances in the cryogenic data collection may revive sub-zero temperature-dependent structural biology, offering insights inaccessible to techniques like NMR and cryo-EM.

## 5. Conclusions

Hydrophobic stabilizers (e.g., Apiezon grease) mitigate crystal instability, enabling temperature-dependent studies across cryo-to-room-temperature regimes. This approach yields raw B-factors critical for resolving harmonic/anharmonic vibrational modes and their coupling to conformational dynamics.

## Data availability

The experimental information and data supporting the findings of this study are available within the paper and the indicated data repository, under PDB ID listed in Table S1 for data collected at 100 K using Apiezon N (9P0Y, 9P0Z, 9P10), Apiezon T (9P56, 9P57, 9P58), lard (9P5B, 9P5D, 9P1W), glycerol (9P59, 9P5A, 9P5E), mineral oil (9P5F, 9P5G, 9P5H), no cryo (9P5J, 9P5K, 9P5L), olive oil (9P5M, 9P5N, 9P5O), PEG 400 (9P5P, 9P5Q, 9P5R), PEG 6,000 (9P5S, 9P5T, 9P5U), silicone oil (9P5V, 9P5W, 9P5X), vaseline (9P5Y, 9P5Z, 9P60), and at 300 K using Apiezon N (9P61, 9P62, 9P63), Apiezon T (9P64, 9P65, 9P66) or vaseline (9P67, 9P68, 9P69). Further information is available from the corresponding authors upon reasonable request.

## Abbreviations

HEWL: hen egg white lysozyme;

## Acknowledgments

We would like to thanks the staff from Manaca beamline and LNBio (Sirius, Centro Nacional de Pesquisa de Energia e Material - CNPEM, Campinas, SP, Brazil) and CENABIO-UFRJ for support.

## Funding

This study was supported by Fundação Carlos Chagas Filho de Amparo à Pesquisa do Estado do Rio de Janeiro (FAPERJ; grants E-26/202.998/2017-BOLSA, E-26/200.833/2021-BOLSA, E-26/010.001434/2019-Tematico, E-26/210.195/2020 and SEI-260003/001207/2023 - APQ1-Tematico to LMTRL), by the Conselho Nacional de Desenvolvimento Científico e Tecnológico (CNPq; grant PQ/311582/2017-6; PQ/313179/2020-4; 311784/2023-2 to LMTRL; INCT Structural Biology and Bioimage), the Financiadora de Estudos e Projetos (Brazilian Funding Authority for Studies and Projects, FINEP; Grant #01.11.0100.00), by the Coordenação de Aperfeiçoamento de Pessoal de Nível Superior (CAPES, Finance Code #001), and by grants from the CNPEM. The funding agencies had no role in the study design, data collection and analysis, or decision to publish or prepare of the manuscript.

## Conflict of interest

The authors declare no financial or intellectual conflicts of interest with the contents of this article.

## Accession Code

Hen egg white lysozyme (UNIPROT ID P00698)

## Author Contributions

**Fernando de Sá Ribeiro** - Data curation; Formal analysis; Investigation; Methodology; Validation; Visualization; Writing - review & editing.

**Luís Maurício T. R. Lima -** Conceptualization; Data curation; Formal analysis; Funding acquisition; Investigation; Methodology; Project administration; Supervision; Validation; Visualization; Roles/Writing - original draft; Writing - review & editing.

## Notes

### Competing Interest Statement

The authors have declared no competing interest.

