## Supporting Material for "Pinpointing protein structures over broad temperature range using hydrophobic protection"

[1] Laboratório de Biotecnologia Farmacêutica (pbiotech), Faculdade de Farmácia, Universidade Federal do Rio de Janeiro, Rio de Janeiro, RJ, 21941-902, Brazil.
[2] Programa de Pós-Graduação em Química Biológica, Universidade Federal do Rio de Janeiro, Rio de Janeiro, RJ, 21941-902, Brazil.
[3] Programa de Pós-Graduação em Ciências Farmacêuticas, Faculdade de Farmácia, Universidade Federal do Rio de Janeiro, Rio de Janeiro, RJ, 21941-902, Brazil.
[4] Programa de Pós-Graduação em Nutrição, Universidade Federal do Rio de Janeiro, Rio de Janeiro, RJ, 21941-902, Brazil.

**Running title:** Broad-range temperature-resolved crystallography

*To whom correspondence should be addressed

**AUTHOR LIST**

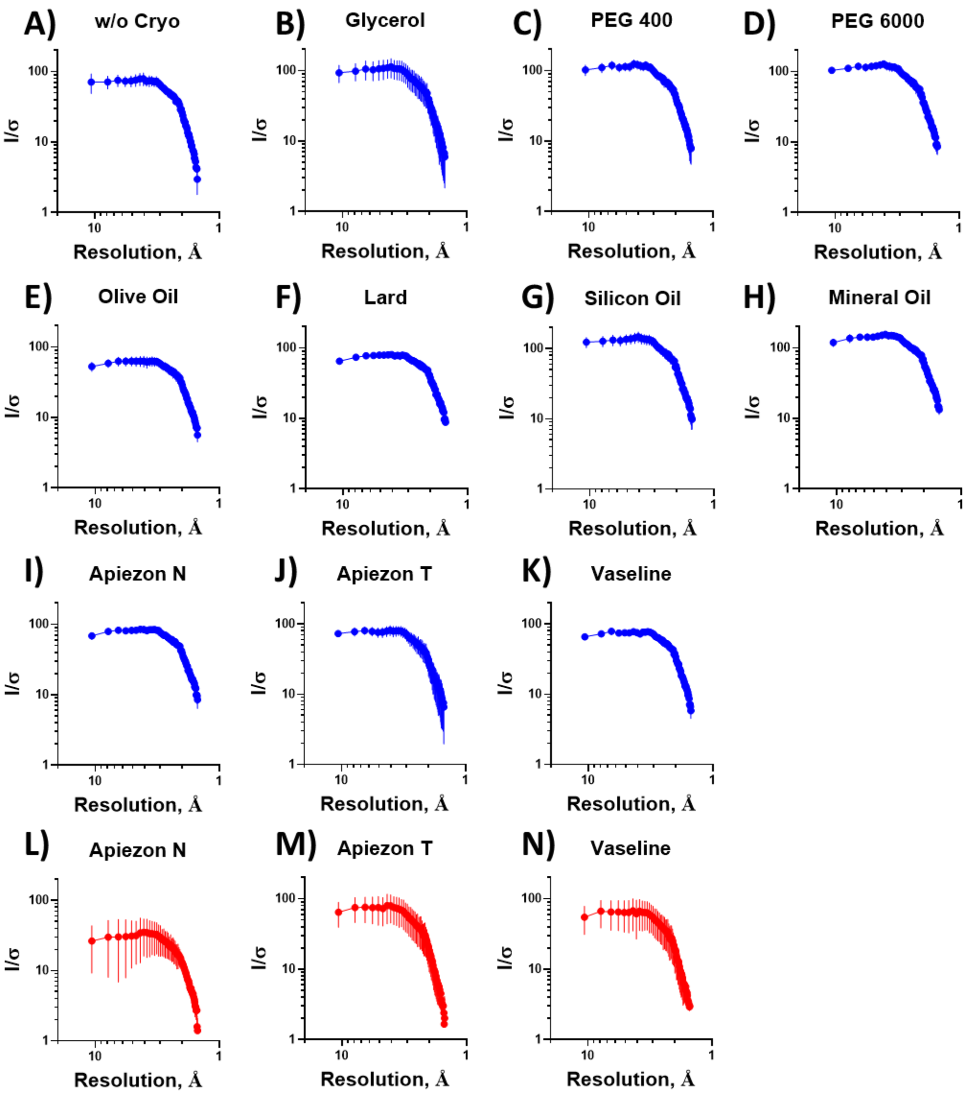


**Figure S1. I/σ distribution by resolution.** Crystals were diffracted using home-source CuKα radiation at 1.5 Å resolution from D8 Venture diffractometer (Bruker AXS Inc.), integrated using Proteum*3* (Bruker AXS Inc.) and scaled using Aimless (CCP4). The Wilson plot is presented for crystals (n=3 per condition) collected at **100 K** (A – K) and **300 K** (L – N). Details in the Experimental Section of the corresponding manuscript.

**Table 1. Crystallographic data**. Crystals collected at home-source and synchrotron were processed and a summary of the statistical data are presented in the corresponding spreadsheet associated to this work, including Temperature, protectant agent, PDB ID, cell parameters, Wilson B-factor, average B-factor, I/σLow, I/σHigh, $CC_{1/2}$, , Completeness, among others.
